# Terraces in Gene Tree Reconciliation-Based Species Tree Inference

**DOI:** 10.1101/2020.04.17.047092

**Authors:** Michael J. Sanderson, Michelle M. McMahon, Mike Steel

## Abstract

Terraces in phylogenetic tree space are sets of trees with identical optimality scores for a given data set, arising from missing data. These were first described for multilocus phylogenetic data sets in the context of maximum parsimony inference and maximum likelihood inference under certain model assumptions. Here we show how the mathematical properties that lead to terraces extend to gene tree - species tree problems in which the gene trees are incomplete. Inference of species trees from either sets of gene family trees subject to duplication and loss, or allele trees subject to incomplete lineage sorting, can exhibit terraces in their solution space. First, we show conditions that lead to a new kind of terrace, which stems from subtree operations that appear in reconciliation problems for incomplete trees. Then we characterize when terraces of both types can occur when the optimality criterion for tree search is based on duplication, loss or deep coalescence scores. Finally, we examine the impact of assumptions about the causes of losses: whether they are due to imperfect sampling or true evolutionary deletion.

---

A long standing and still largely dominant paradigm in phylogenetic tree inference is based on optimization of some score derived from data over candidate tree solutions. In addition to familiar maximum parsimony, maximum likelihood, and (certain) distance-based methods like minimum evolution and Fitch-Margoliash (Felsenstein 2004), all commonly used to infer a tree from a sequence alignment, optimization methods also are employed in a diverse set of methods aimed at solving other tree inference problems, such as supertree construction, gene tree reconciliation, species tree inference using likelihood or pseudo-likelihood, and network reconstruction. Computational obstacles in optimization include the problem of multiple optima and regions where the solution space is flat, which can both impede algorithms to find optima and make circumscription of solutions more complex. One contributor to this problem in phylogenetics is missing data, and a particularly direct example of this is the phenomenon of “terraces”—regions of tree space having identical optimality scores purely due to certain patterns of missing data among the taxa sampled (Sanderson et al. 2011, 2015).

The properties of terraces have been elucidated mostly in the context of large multilocus data sets, where the pattern of missing data can be described by the “taxon coverage” of data—which loci are sampled for which taxa. If a tree is inferred for a concatenated multilocus alignment by maximum parsimony, or by maximum likelihood with certain model assumptions, the pattern of taxon coverage alone can be used to infer the number and sizes of terraces having identical optimality scores. Surveys of empirical studies indicate that terraces can be astronomically large in large trees (Dobrin et al. 2018), and they can degrade other aspects of phylogenetic inference aside from tree search, such as estimation of confidence intervals (Sanderson et al. 2015). They are likely to have an impact on comparative methods and other studies that employ statistical inference methods that depend on accurate evaluation of the confidence set of trees.

The conditions required for terraces to be observed in phylogenetic inference problems are quite general (Steel 2016). In this paper we describe their impact on a large additional set of phylogenetic problems involving building species trees from gene trees using gene tree reconciliation methods. This includes methods that have been used to infer species trees from gene families that have undergone a history of gene duplication and deletion, as well as methods that have been used to infer species tree from allele trees in which population genetic processes lead to deep coalescence events. This area is undergoing active development in the phylogenomics community and our results pertain most directly to discrete parsimony-like methods arising from the reconciliation framework, but potential connections to model based approaches may exisit by analogy to previous findings with maximum likelihood in concatenation methods (Sanderson et al. 2011). We find that terraces are not only expected in this new context, but that an additional *type* of terraces can be seen in certain versions of reconciliation problems.

## Trees, Subtrees, and Display

We assume all trees, *T*, are rooted, binary, and have edges, *E*(*T*) and nodes, *V* (*T*), and root node, *r*(*T*) ∈ *V* (*T*). Nodes are partitioned into the set of internal nodes, 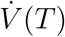 (having outdegree two or more), and the set of leaves, *L*(*T*) (having indegree one), each of which is labeled by a distinct element of the leaf set, *X*. If node *v*′ is an ancestor of *v*, we write *v*′ *> v*. The set of leaf labels descended from *v* is *C*_*T*_ (*v*). The node of the most recent common ancestor of a set of leaves, *A*, is MRCA_*T*_ (*A*).

Given a rooted binary tree, *T* with leaf label set, *X*, and with *X*_*g*_ ⊆ *X*, we define two kinds of subtrees of *T* having leaf label set, *X*_*g*_ (Fig. 1):

1. The “homomorphic subtree”, 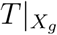, is the smallest subtree of *T* having leaf label set *X*_*g*_ and suppressing any interior nodes with outdegree one (“unary” nodes). When the context is clear, we abbreviate this to *T* |_*g*_.
2. The “restriction subtree”, 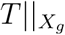, is the smallest subtree having leaf label set *X*_*g*_ but keeping any unary nodes. When the context is clear, we abbreviate this to *T*∥_*g*_.

**Fig. 1.**
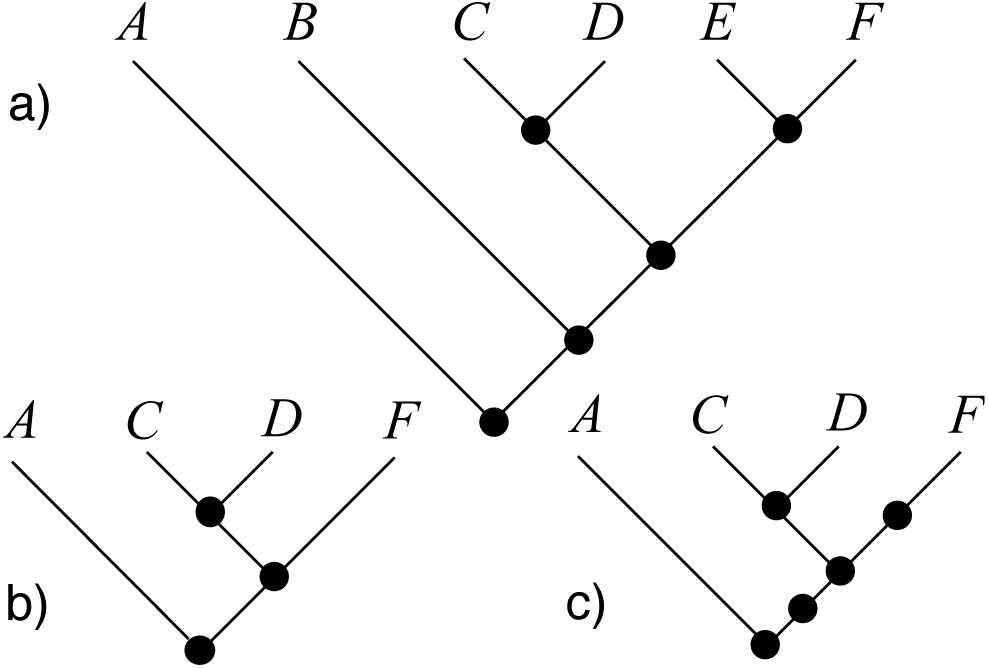
Illustration of two subtree operations, retaining leaf label set *X*_*g*_ = {*A, C, D, F*}: a) Original species tree, *T* ; b) homomorphic subtree, 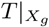; c) restriction subtree, 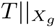.

A tree *T displays* a binary subtree *g* having leaf label set *X*_*g*_ ⊆ *X* if 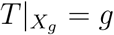 (Steel 2016). [If *g* is not binary then this definition can be extended to allow *T* |_*g*_ to be a refinement (resolution) of *g*.] This is the conventional definition of “display”, but it can be generalized to use the restriction subtree when appropriate (see below).

## Spans and Terraces

In general, given a sequence of rooted binary trees, 𝒯 = (*T*_1_, …, *T*_*k*_), with leaf label sets *X*_*i*_ (*X* = ⋃_*i*_*X*_*i*_), define the *span*, ⟨𝒯⟩, as the set of all rooted binary trees having label set *X* that display every tree in 𝒯 (Fig. 2). The span may be empty.

**Fig. 2.**
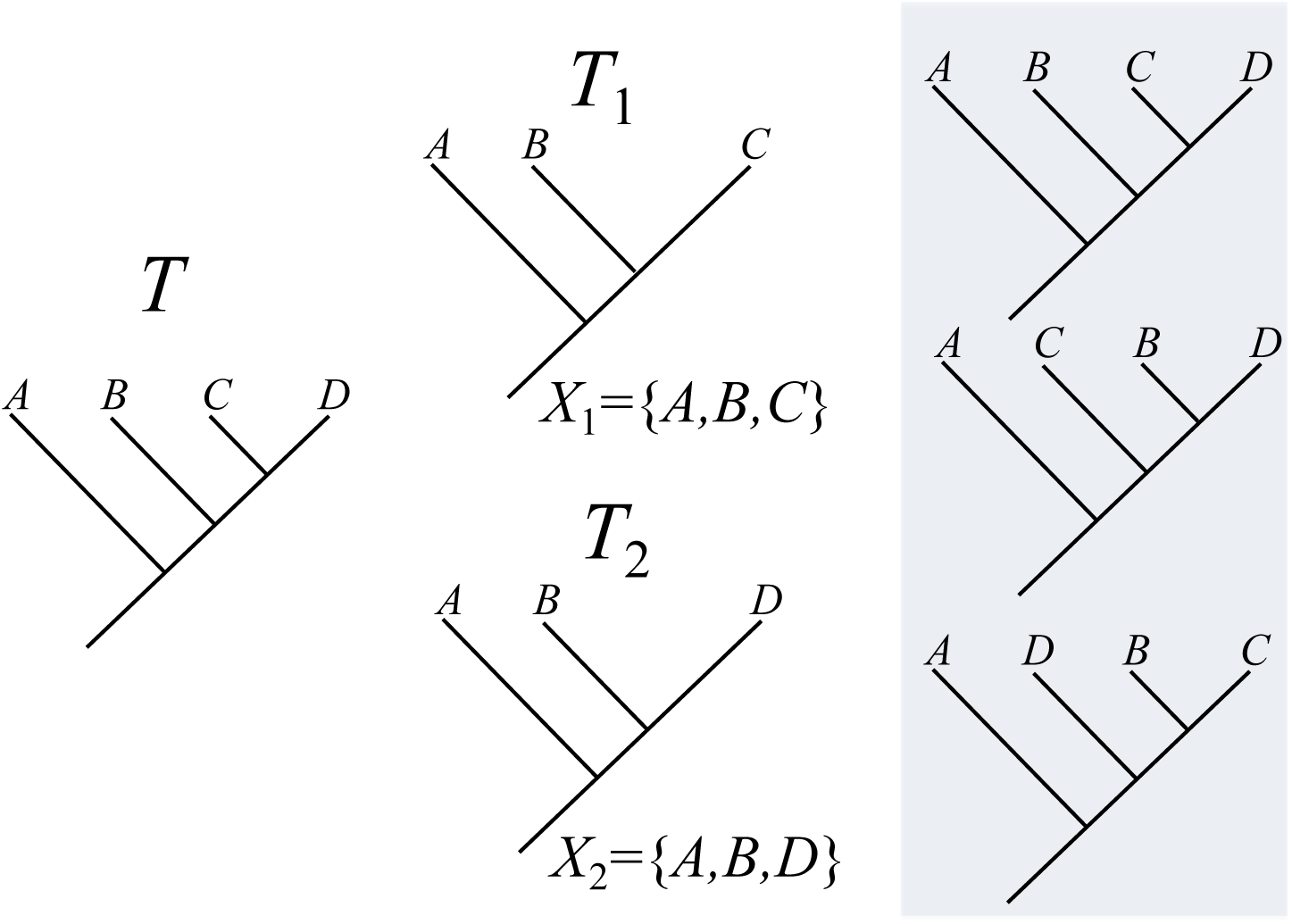
Span of a set of subtrees. Original rooted parent tree is *T* having leaf label set, *X* = {*A, B, C, D*}. Subsets of label sets, *X*_1_ and *X*_2_, induce a sequence of subtrees, 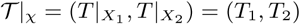. Each of the trees on *X* in the shaded box displays both *T*_1_ and *T*_2_ and this set of three trees is the *span* of 𝒯|_*χ*_, ⟨𝒯|_*χ*_⟩.

Consider the special case of a rooted binary *parent tree, T*, with leaf label set, *X*, along with a sequence of leaf label subsets, *χ* = (*X*_1_, …, *X*_*k*_), *X*_*i*_ ⊆ *X*. Label set *X*_*i*_ may be thought of as the set of leaves present in a subtree of *T* or as the set of leaves of *T* that have some kind of data present, say the sequence of the *i*th gene in a multigene multiple sequence alignment. Let 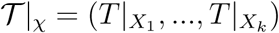 then ⟨𝒯|_*χ*_⟩ is the span of this sequence of subtrees derived from the parent tree (Fig. 2). Clearly the size of this span is at least one since it contains *T*.

A *terrace* is a span, ⟨𝒯|_*χ*_⟩, in which all the trees have the same *score*. The properties of this score determine whether this is a possibility. Given some cost function, *c*(*S, g*) for a tree, *T*, with leaf label set, *X*, based on some data associated with “gene”, *g*, valid even if only a subset of leaves, *X*_*g*_ ⊆ *X*, have data present; and given some score function, *s*(*c*(*T, g*_1_), …, *c*(*T, g*_*m*_)), that combines these costs across a set of genes, 𝒢 = {*g*_1_, …, *g*_*m*_}, the following two properties of *c* and *s* are sufficient to cause all trees in the span to have the same score (Sanderson et al. 2011; Steel 2016):

### Condition 1.

The cost function is the same on the full tree as it is on the subtree pruned to those taxa that have data.

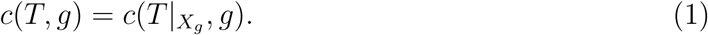

### Condition 2.

The score function, *s*, is a linear sum of costs for each gene:

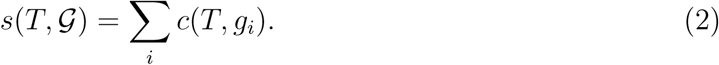

### Theorem 1 (Terraces—Type I).

*If the score function, s, satisfies the two conditions described above, then every tree in the span*, ⟨𝒯|_*χ*_⟩, *has the score, s*(*T*, 𝒢).

*Proof.* Following Steel (2016), if *T* ′ ∈ ⟨𝒯|_*χ*_⟩, Then 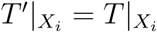 for *i* = 1, …, *k*, so

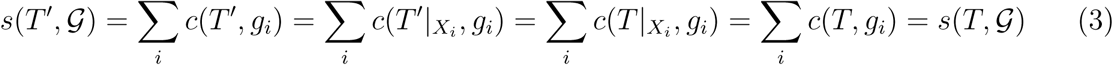

□

This is most interesting if |⟨𝒯⟩| ≫ 1. In fact, the size of terraces can be astronomically large for some data sets for large parent trees (Sanderson et al. 2011; Dobrin et al. 2018) when there is a sizable amount of missing data.

In Sanderson et al. (2011, 2015) and Steel (2016) the data set associated with gene *g* was taken to be a multiple sequence alignment over some block of sites, and the scores considered included maximum parsimony and maximum likelihood scores. The exact circumscription of “blocks” is relevant, but for simplicity we conceptualize it as data associated with a single gene in the genome (see Sanderson et al. 2015).

### Terraces—Type II

We now show that terraces arise when using the restriction subtree operation, which will be relevant for discussion of gene tree reconciliation below.

Define a “restriction span” by (i) using the restriction subtree in the above definition of “display”, and then (ii) letting 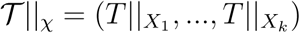. The resulting span can be labeled ⟪𝒯∥_*χ*_⟫ (see example in Fig. 3). [The double angle brackets reinforce the idea that the definition of “display” has changed]

**Fig. 3.**
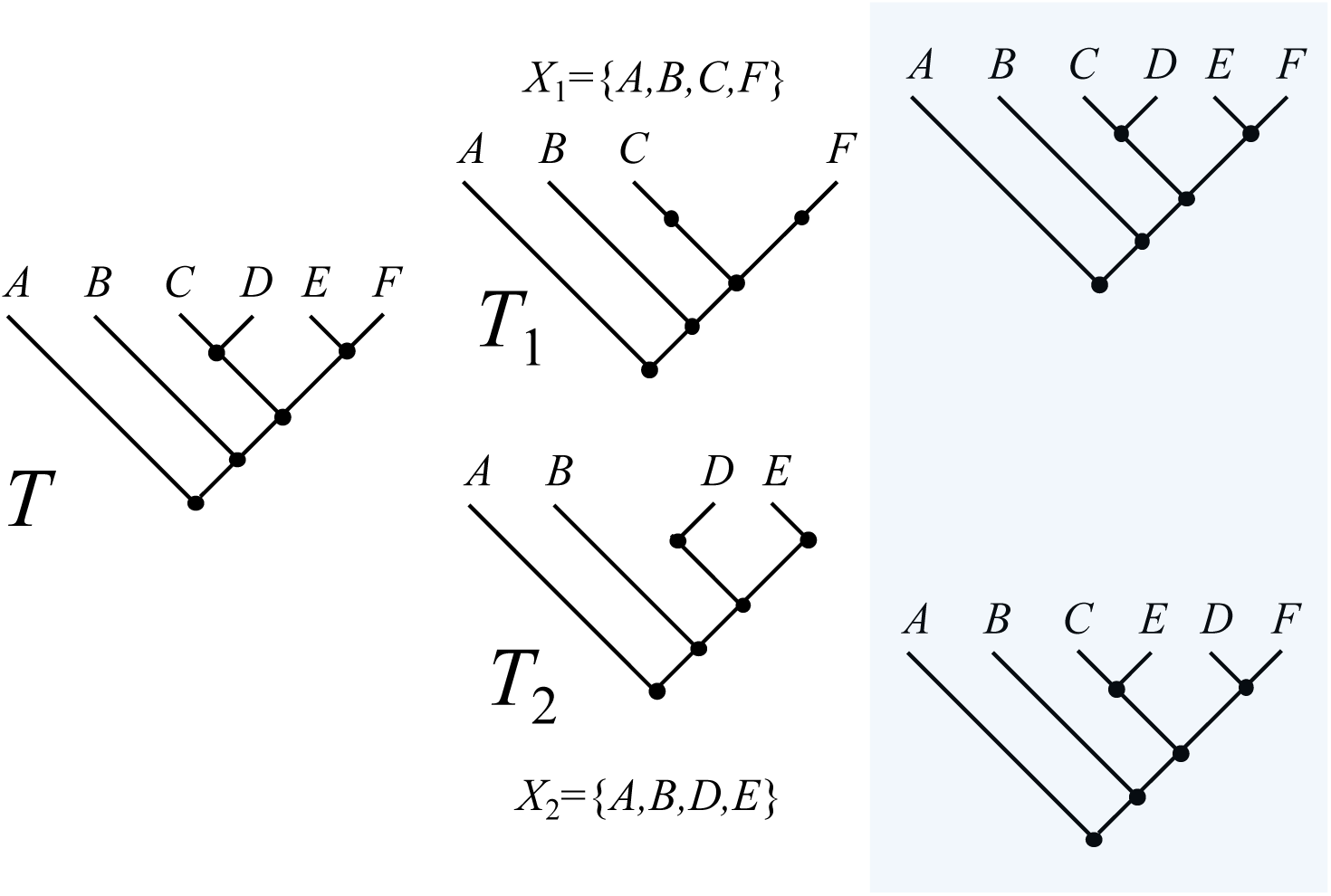
Restriction span of a set of subtrees. Original rooted parent tree is *T* having leaf label set, *X* = {*A, B, C, D, E, F*}. Subsets of label sets, *X*_1_, *X*_2_ ⊆ *X*, induce “restriction” subtrees (Fig. 1), 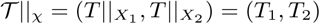. Each of the trees in the shaded box displays both *T*_1_ and *T*_2_, and this set of two trees is the *restriction span*, ⟪𝒯∥_*χ*_⟫.

#### Theorem 2 (Terraces—Type II).

*If the score function, s, satisfies the second condition required for Theorem 1 (linear sum of costs), and the first condition is modified so that* 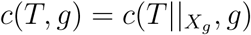, *then every tree in the restriction span*, ⟪𝒯|_*χ*_⟫, *has the score, s*(*T*, 𝒢).

*Proof.* The proof follows easily by substituting the restriction span and restriction subtree operations in the proof for Theorem 1 □

The example in Fig. 3 shows that there can be more than one tree on this kind of terrace, as with the first type of terrace. The elements of ⟪𝒯∥_*χ*_⟫ and ⟨𝒯|_*χ*_⟩ are not necessarily the same. For example, in Fig. 3 there are 15 trees in ⟨𝒯|_*χ*_⟩ but only two in ⟪𝒯∥_*χ*_⟫. In general, the former set of trees contains the latter, as we now state:

#### Theorem 3.

*For any rooted binary parent tree, T, having leaf label set X, and given label sets χ* = (*X*_1_, *…, X*_*k*_), *X*_*i*_ ⊆ *X*,

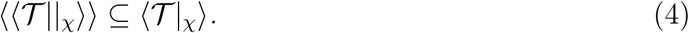

*Proof.* Suppose that 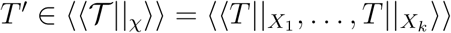 Then 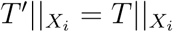 for all *i* = 1, *…, k* and so 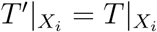 for each *i* (since restriction display implies ordinary display) and thus 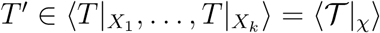 □.

In parallel with the terminology for terraces, we refer to these two spans as “Type I” and “Type II” respectively.

## Species Tree Inference by Reconciliation

Different regions of the genome can have different phylogenetic histories, and these “gene trees” can also differ from the species tree in which they are imbedded (Bravo et al. 2019; Liu et al. 2019). The framework of gene tree “reconciliation” provides an analytically rich and empirically powerful toolkit to understand this mosaic of histories and to infer species trees from discordant gene trees (Goodman et al. 1979; Page 1994). Here we extend our previous results on terraces to the reconciliation setting by discussing conditions under which reconciliation-based score functions lead to terraces when there are missing leaves in the gene trees.

Reconciliation algorithms are based on costs that reflect (at least) four kinds of explicitly *evolutionary* processes in the phylogenetic history of a gene tree imbedded in a species tree, “speciation”, “duplication”, “deletion”, and “deep coalescence”. In addition, costs may reflect some aspect of *sampling* of leaves (Page 1994; Page and Charleston 1997). Additional evolutionary events, such as lateral transfer, have been studied to a lesser extent (Bansal et al. 2012) but are not considered here. Assume there is a rooted binary species tree, *S*, leaf labeled by ℒ_*S*_, and an imbedded rooted binary gene tree, *g*, labeled by ℒ_*g*_ ⊆ ℒ_*S*_. In general, all reconciliation cost functions can be written as *c*(*S, g*), and these are generally combined additively across gene trees, so the score function *s*(*T*, 𝒢) discussed above extends easily in principle, even though the underlying sequence data are no longer relevant.

Define a *reconciliation* of *S* and *g*, loosely, as an annotation of *g* that indicates the locations of speciation events, duplications and losses. [More formal definitions include defining a *reconciled tree* as an extension of *g* made by inserting “lost” subtrees so that the resulting tree is consistent with speciation and duplication alone (Chauve and El-Mabrouk 2009).]

Given a set of gene trees, 𝒢, the *species tree inference problem* is then to find a species tree, *Ŝ*, that minimizes

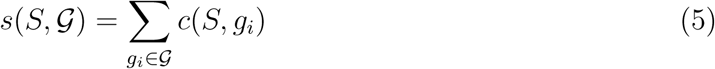

over all *S*, for various choices of the cost function, *c*(*S, g*_*i*_).

## Incompleteness, missing data, absence, deletion and loss

Let ℒ_*S\g*_ be the set of leaf labels of *S* missing from *g*. If ℒ_*S\g*_ ≠ Ø, then *g* is *incomplete*. The concept of “missing data” in the original notion of terraces equates to incompleteness in one or more gene trees, and this can lead to terraces in the species tree inference problem.

However, distinguishing between evolutionary events and sampling processes can be confounded in incomplete gene trees. In a gene tree having no leaves from species *x*, were these genes evolutionarily deleted, or was *x* never sampled? For *x* ∈ ℒ_*S\g*_, by “incompleteness due to deletion”, we mean that the absence of *x* from *g* is considered positive evidence that no such leaves exist in *g* (*x* has been well studied and its absence is meaningful)—absence implies deletion. By “incompleteness due to sampling”, we mean *x* has not been studied, and we make no claim a priori about whether further study of *x* would reveal it has leaves in *g*—absence implies no more than failure to sample or bad lab technique (Bayzid and Warnow 2018).

Page and Charleston (1997) pointed out that the loss cost could influence species tree inference under reconciliation and thereby raised the issue of how its interpretation could matter. Bayzid and Warnow (2018) showed, in fact, that the correct loss score of a gene tree reconciled with a species tree depends directly on this assumption about the meaning of loss. We therefore qualify the term “loss” by whichever mechanism is assumed to generate it.

## Reconciliation Costs, Incompleteness and Spans

When *g* is complete, it is well known that an optimal reconciliation exists that minimizes the number of duplications, losses, and duplications plus losses (Chauve and El-Mabrouk 2009). The solution can be found using the *MRCA-mapping*, ℳ, from node *u* in *g* to a node in *S*: ℳ_*S*_(*u*) = MRCA_*S*_(*C*_*g*_(*u*)), which identifies the lowest node on the species tree that can have the subtree of *g* rooted at *v* imbedded within it (Fig. 4). Briefly, in this optimal reconciliation, a node *u* on *g* is annotated as a duplication if ℳ_*S*_(*u*) maps to the same node of *S* that one of the child nodes of *u* does. The duplication cost, *c*_*dup*_, is the number of such duplications. Determining the location of, and the number of losses, *c*_*loss*_, is just slightly more complicated, and depends on lengths of paths mapped onto *S* from *g* using ℳ_*S*_(*v*). The duplication plus loss cost is then *c*_*D*+*L*_ = *c*_*dup*_ + *c*_*loss*_. All of these computations have been described in detail elsewhere (e.g. Zhang 2011; Bayzid and Warnow 2018) and we omit them here.

**Fig. 4.**
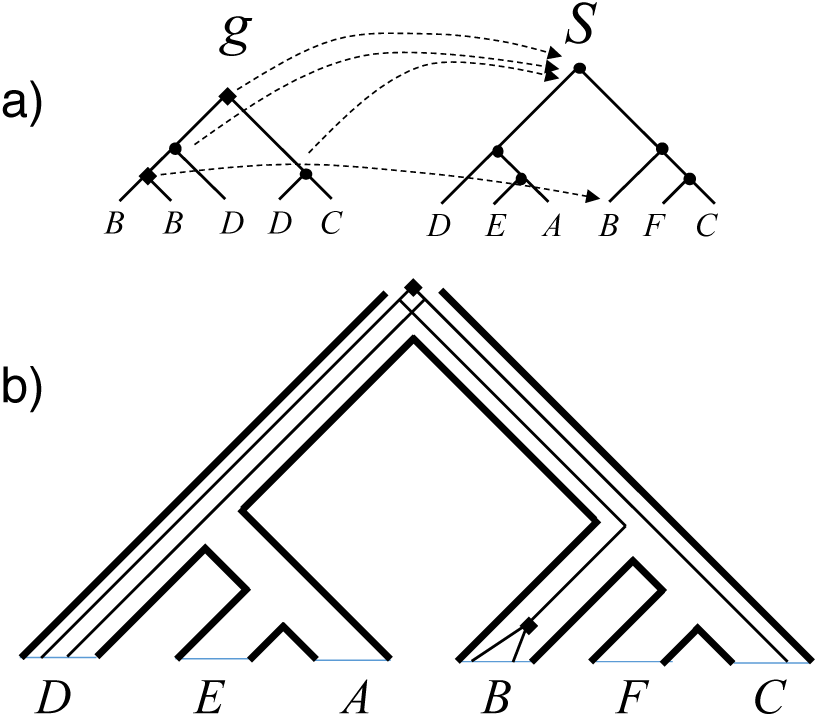
Reconciliation of a gene tree, *g*, and species tree, *S*. Note the set of “missing leaves” on *g*, ℒ_*S\g*_= {*A, E, F*}. a) The MRCA mapping, ℳ, is shown for all internal nodes of *g*. Nodes annotated as duplications on *g* indicated by diamonds; nodes annotated as speciation indicated by dots. b) Imbedding of *g* in *S*. For clarity, only the duplication nodes on *g* are shown as annotated.

There are multiple formulations of the deep coalescence cost (Zhang 2011; Steel 2016), *c*_*DC*_, which is the number of “extra edges” of the gene tree imbedded within the species tree for this reconciliation. Any edge *e* = (*u, v*) ∈ *E*(*g*) is imbedded in *k*_*S*_(*e*) ⩾ 0 edges of *S* where *k*_*S*_(*e*) is the number of edges in the path from ℳ_*S*_(*u*) to ℳ_*S*_(*v*), so the number of extra edges is,

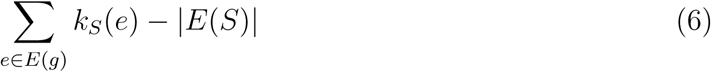

since there is minimally one edge in each edge of *E*(*S*).

The computation of these reconciliation costs for incomplete gene trees depends in part on the interpretation of incompleteness.

### Incompleteness due to sampling

The trees *g* and *S*|_*g*_ have the same label sets and standard reconciliation algorithms can be used to compute *c*(*S*|_*g*_, *g*), but how do we compute *c*(*S, g*)? If the absence of a gene tree leaf for some species taxon is assumed to arise because of failure to sample it, we compute the cost, *c*(*S, g*), by “completing” the gene tree so it has the same label set as the species tree (Bayzid and Warnow 2018). A *completion, g*′, is the tree obtained from *g* by adding subtree(s) having all the leaves, *x* ∈ ℒ_*S\g*_. An *optimal completion, g*\*, with respect to some cost function, *c*, is the completion that minimizes *c* over all *g*′. For example, the optimal completion with respect to the duplication cost would be that gene tree, *g*\*, that has smallest number of duplications among all possible completions. The optimal completion is a guess about unsampled leaves that disturbs the cost the least. From here on, when discussing optimal completions for incomplete gene trees, we use as shorthand, *c*(*S, g*), instead of *c*(*S, g*\*(*g*)).

Bayzid and Warnow (2018) show that there are optimal completions for the duplication and loss scores, which have the same duplication and loss scores as that for *c*(*S*|_*g*_, *g*), so that:

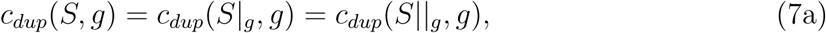

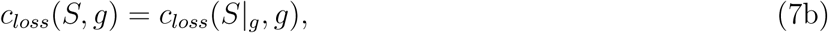

which implies that the duplication plus loss cost, *c*_*D*+*L*_ can be obtained simply by adding these together. Notice the subtree operation for the loss cost is the homomorphic subtree operation.

This can be shown by the following argument. Each unary node, *u*, of *S*∥_*g*_ corresponds to a binary node on *S* that has one pendant subtree, *t*, the leaves of which are all in ℒ_*S\g*_. (i.e., missing). If there are *l* edges of *g* passing through *u*, then we attach *l* replicate instances of *t* along their respective edges of *g* at new nodes *v*_*i*_, *i* = 1, …, *l*, of *g*. For each *v*_*i*_, the mapping, ℳ(*v*_*i*_) must be to a different node in *S* than either the parent of *v*_*i*_ maps to, or any of its children map to, because the leaves of *t* are in ℒ_*S\g*_. Thus there are no additional duplications inferred by this completion. This argument holds equally for both subtree operations, so *c*_*dup*_(*S*|_*g*_, *g*) = *c*_*dup*_(*S*∥_*g*_, *g*).

This completion also avoids adding *l* losses, by “imputing” just the right leaves to be present as subtrees of *g* (Fig. 5). The loss cost is computed on *S*|_*g*_ but not *S*∥_*g*_. The intuition for this is that adding these “ghost” subtrees, *t*, at edges of *g* passing through *u* allows us to pretend, with respect to losses, that node *u* was never there.

**Fig. 5.**
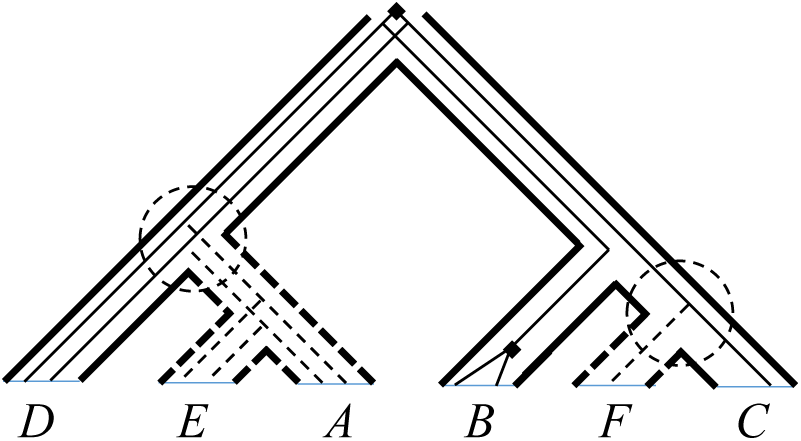
Optimal completion under the duplication-loss costs. Trees are the same as in Fig. 4. Dashed border for some species tree edges indicates parts of *S* that are missing from *g* because of absence of leaves *A, E, F*. Dashed circles are unary nodes on the restriction subtree, *S*∥_*g*_. Dashed edges of *g* represent the optimal completion of *g* that allows leaves of *g* to be “present” at leaves of *S*, while adding as few duplications and losses as possible to the reconciliation. Here, no duplications or losses are added by this completion. Note that for the unary node at left, two edges of *g* traverse it and two subtrees are added, whereas for the unary node at right having one imbedded edge, only one subtree need be added.

For the deep coalescence cost when incompleteness is assumed due to sampling we can also use the optimal completion approach (Bayzid and Warnow 2012, 2018)(Fig. 6). First note that the argument above regarding duplications remains true and no new duplications need be added by this completion. Now at each unary node, *u*, of *S*∥*g*, however, we attach only one instance of *t* to the *l* ⩾ 1 edges of *g* passing through *u*. The remaining *l* − 1 edges split at *u* but immediately are lost (and the interpretation of loss does not matter). This guarantees that at most one edge of *g* will be present in *t* and therefore the DC cost will not increase relative to what it was with respect to *S*∥*g*, which means

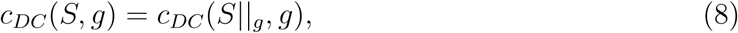

which, from Eq. 6, means

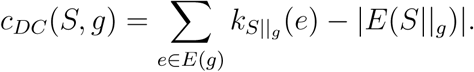

**Fig. 6.**
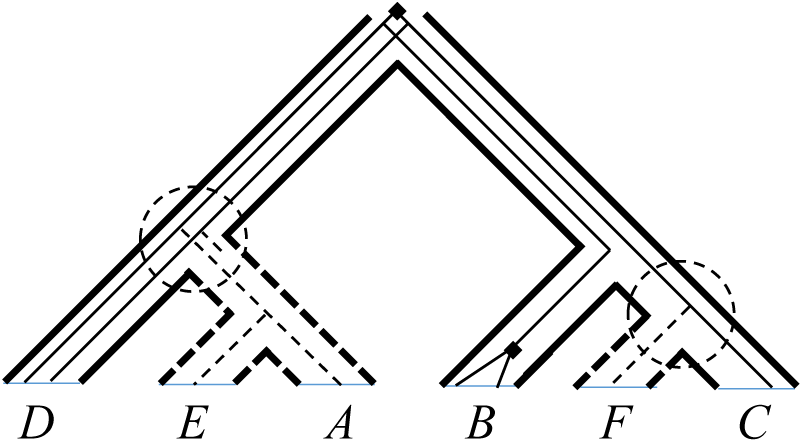
Optimal completion under the deep coalescence cost. Layout is the same as in Fig. 5. Here the optimal completion must add as few “extra edges” as possible (i.e., over and above one edge of *g* per edge of *S*), regardless of how many losses are then required. Note the loss of one edge of *g* immediately after its split within the unary node of *S* at lower left. Contrast with Fig. 5.

Note that this is not necessarily an optimal completion for *losses*, as it is fine to add losses in order to keep deep coalescences to a minimum.

### Incompleteness due to deletion

Here the absence of a gene tree leaf is assumed to be caused by evolutionary deletion of that gene somewhere in the tree. With this assumption, we still have

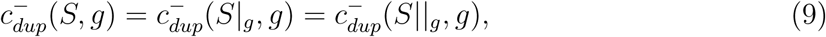

(Chauve et al. 2008; Górecki and Tiuryn 2006; Bayzid and Warnow 2018), where we add the superscript (^−^) to *c* to refer to the deletion interpretation of absence. Duplications are unaffected by the choice of subtree operations, because ℳ does not map nodes of *g* to unary nodes of *S*, and these nodes are the only difference between *S*|_*g*_ and *S*∥_*g*_.

But the same is not true for the loss cost (Bayzid and Warnow 2018). The correct count of losses when incompleteness is due to deletion is computed by substituting *S* for *S*|_*g*_ in Theorem 3 of Bayzid and Warnow (2018). This computation uses lengths of paths on *S* that run from some node ℳ_*S*_(*u*) to ℳ_*S*_(*v*), where *u, v* is a child and parent node on *g*. Together, all paths in *S* of this type comprise the tree, *S*∥_*g*_, which means

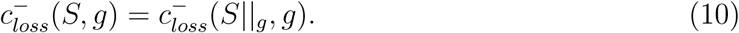

So, unlike the duplication cost, the loss cost is not necessarily the same under the two interpretations of incompleteness. In fact, they differ exactly by which kind of subtree operation is appropriate. Bayzid and Warnow (2018, Thm. 4) give an example in which these different interpretations of loss lead to different optimal species trees when using the loss cost.

Bayzid and Warnow (2018) also discussed the special case in which missing leaves are distributed on the gene tree in a way that might imply a gene is absent from the root of the species tree (Fig. 7). This is not a problem when incompleteness is assumed to arise from sampling, but if it is assumed to arise from deletion, it is necessary to make an assumption about whether genes are present at the root. Let *u*\* = ℳ_*S*_(*r*(*g*)) be the location on *S* of the root of *g*. If *r*(*S*) *> u*\*, should we assume the gene is present at *r*(*S*)? In the above treatment, the definition of *S*∥_*g*_ solves the loss problem under the assumption that a gene is unambiguously present only in the subtree of *S* having *u*\* as its root. To make the stronger assumption that it is present at *r*(*S*), we can define yet another subtree operator, 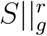, which is the smallest subtree of *S* containing *r*(*S*) and the leaf labels of *g*. The original restriction operator, *S*∥_*g*_, can be interpreted in this problematic boundary case to imply losses are due to deletion in the subtree of *S* having *u*\* as its root, and losses are due to sampling elsewhere. We do not pursue this further here, but see Bayzid and Warnow (2018).

**Fig. 7.**
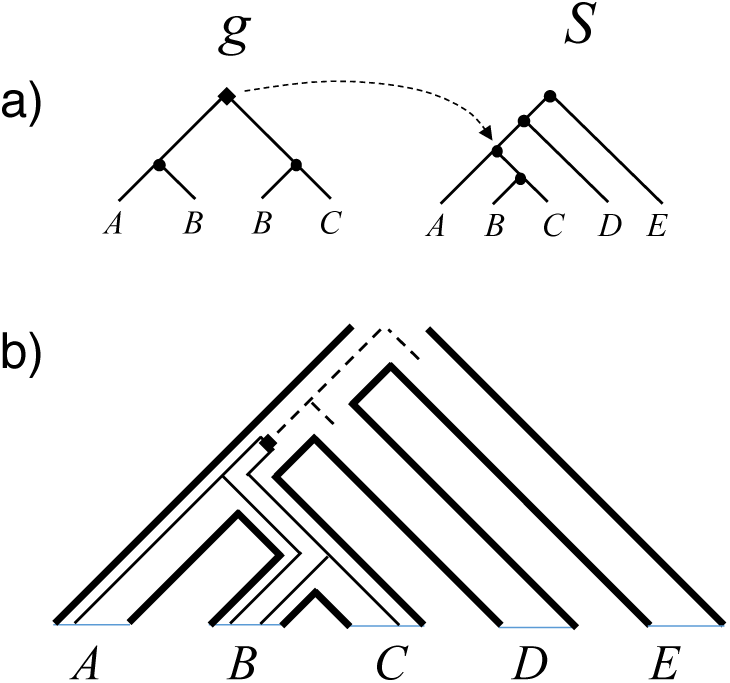
Special case in which presence of gene at the root of the species tree is ambiguous for incompleteness due to deletion. a) Incomplete gene tree and species tree with MRCA mapping (ℳ) from root of *g* shown. b) Imbedding of *g* in *S* leaving uncertainty toward the root of *S*. Dashed lines represent presence of gene if it is forced present at root of *S* and location of two losses entailed as a consequence.

The deep coalescence cost when incompleteness is assumed to arise from deletions parallels the case for incompleteness due to sampling. On *S*∥*g*, any unary node, *u*, has *l* ⩾ 1 edges of *g* passing through it. To explain the missing leaves present in the pendant subtree, *t*, of *u* that is present on *S* but not *S*∥*g*, each of these *l* lineages must branch at *u* and end in a deletion somewhere in *t*. Moreover, at most one edge of *g* can persist in *t*, else there is a deep coalescence and the DC cost is increased. This lineage must end in a deletion before reaching a leaf node of *S*, and in fact it can be deleted immediately after splitting without changing the DC score, because if an edge of *S* has no imbedded edges of *g*, its DC score is still zero. In this respect the final form of *g* can differ from the optimal completion under sampling, but the net result is the same. All missing leaves can be accounted for without adding to the DC cost (though perhaps adding to the number of losses), and thus the DC cost is the same as for incompleteness due to sampling:

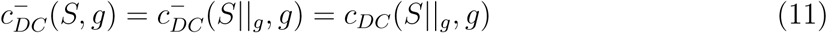

## Reconciliation and Terraces

Based on the reconciliation costs for incomplete gene trees in Eqs. 7-11 above and Theorem 2, we can summarize which combinations of costs and interpretations of losses can lead to terraces (Table 1). First, recall that, from Theorem 3, for a given input, the Type II span of trees is a subset of the Type I span. This implies that if a given reconciliation setting induces Type I terraces (so that scores are the same for all trees in its span), then all trees in the corresponding Type II span will also have the same score and be a Type II terrace. The reverse is not necessarily true.

**Table 1.** Reconciliation Costs and Possible Terrace Types

| Cost/Interpretation of loss | Sufficient Conditions for Terraces of Type |
| --- | --- |
| Duplication/Sampling | I,II |
| Duplication/Deletion | I,II |
| Loss/Sampling/ | I,II |
| Loss/Deletion | II |
| D+L/Sampling | I,II |
| D+L/Deletion | II |
| DC/Sampling | II |
| DC/Deletion | II |

In view of this, all four of the reconciliation costs can lead to Type II terraces and some, in addition, lead to Type I terraces. The duplication cost can lead to both kinds of terraces regardless of the interpretation of incompleteness. In contrast, the DC cost can only lead to Type II terraces (also regardless of the interpretation of incompleteness). For the loss cost, the nature of the assumption about incompleteness determines which type of terrace is possible.

## Discussion

Terraces in tree space can have several deleterious impacts on phylogenetic inference (reviewed in Sanderson et al. 2015), and the basic finding that terraces can arise in tree inference problems under very general conditions (Steel 2016) suggests some care is warranted. Loosely speaking, these conditions involve missing data and an optimality score that is decomposable into additive contributions from different subsets of the data. Originally, terraces were shown to arise in multilocus sequence alignments using maximum parsimony or maximum likelihood optimization scores (Sanderson et al. 2011). In the latter, certain subsets of models lead to terraces and others do not (Sanderson et al. 2011, 2015). Theory (Sanderson et al. 2010) predicted that terraces in these contexts are most likely for data sets with many taxa and few loci. A recent meta-analysis (Dobrin et al. 2018) confirms this expectation and indicates the number of trees on terraces can be very large.

It is no surprise that other classes of data sets and optimality criteria used in phylogenetics that also give rise to terraces. In this paper we explored how terraces can arise in gene tree reconciliation approaches to species tree inference. Here the gene trees take the place of the multiple sequence alignments of individual loci studied in our earlier work (Sanderson et al. 2011), and the optimality scores are additive functions of reconciliation costs that depend on the processes assumed to connect gene trees and species trees. These vary depending on whether processes of gene duplication, loss or incomplete lineage sorting are assumed to occur.

In considering these divergent biological processes, we found a variant of our original phylogenetic terrace that is relevant to some but not all of these processes. For example, under the deep coalescence score, there can be terraces of the second type, but not of the original type described in Sanderson et al. (2011). The composition of trees on a terrace—the “span”—is determined by the pattern of missing data in the input. The span of the trees on a terrace for the deep coalescence score is a subset of the span for the original type of terrace (which would be a terrace for a different optimality score), and therefore the size of this kind of terrace is always less than or equal to terraces of the first type. This implies that the “problem” of terraces should be less for the deep coalescence score than the duplication score, for example.

The phylogenomics literature contains relatively few studies that infer species trees from gene trees subject to duplication and loss (Sanderson and McMahon 2007; Burleigh et al. 2011), compared to those that infer species trees in the context of deep coalescence (e.g. Copetti et al. 2017). For the latter, however, there are a variety of model based and discrete algorithm approaches to inference (Liu et al. 2019), whereas our results are most directly relevant to the simple method of minimizing the deep coalescence score (Maddison 1997; Ma et al. 2001; Zhang 2011; Nakhleh 2013). However, we expect that the general idea of terraces will likely extend to certain model based methods of species tree inference by analogy with how it extends to likelihood based inference from multi-locus sequence alignments (Sanderson et al. 2011, 2015). It remains to be seen whether the ultimate impact of this on species tree inference methods is as dramatic as it is for many large but sparse multiple sequence alignments (Dobrin et al. 2018).

